# Impact of modification of envelope proteins on the mechanical properties of HIV virus-like particles

**DOI:** 10.1101/2025.10.15.682533

**Authors:** Elizabeth Kruse, Michiel van Diepen, Rosamund Chapman, Etienne Horn, Tamer Abdalrahman, Anna-Lise Williamson, Edward P. Rybicki, Wouter H. Roos, Thomas Franz

**Affiliations:** Biomedical Engineering Research Centre, Division of Biomedical Engineering, Department of Human Biology, University of Cape Town, South Africa; Institute of Infectious Disease and Molecular Medicine, Faculty of Health Science, University of Cape Town, Cape Town 7925, South Africa; Division of Medical Virology, Department of Pathology, University of Cape Town, Cape Town 7925, South Africa; Architecture, Construction and Engineering, Berlin International University of Applied Sciences, 10247 Berlin, Germany; Biopharming Research Unit, Department of Molecular and Cell Biology, University of Cape Town, Cape Town 7701, South Africa; Zernike Institute for Advanced Materials, Rijksuniversiteit Groningen, Nijenborgh 4, 9747 AG Groningen, the Netherlands; Bioengineering Science Research Group, Faculty of Engineering and Physical Sciences, University of Southampton, UK

**Keywords:** Virus-like particle, Human immunodeficiency virus, Finite element modelling, Virion mechanics, Elastic modulus, Stiffness

## Abstract

The mechanical interactions between virus-like particles and host cells may offer targets for new viral treatments and vaccines with modes of action that are independent of the immune system. The physical properties of structures involved govern the particle-cell interactions. While the mechanical properties of virions and mammalian cells have been widely studied, data on virus-like particles are limited. This study aimed to determine the mechanical and morphological properties of HIV-1 virus-like particles with different envelopes. Three HIV-like particles, i.e. Gag^M^ + gp150, Gag^M^ + gp140HA_2_tr, and Gag^M^ + gp120HA_2_, were produced by combining the same Gag protein shell with different trimeric glycoprotein envelopes. The particles’ spring constant, breaking force, and dimensions were determined using atomic force microscopy, and the elastic modulus was quantified using finite element analysis. Spring constant, elastic modulus, and breaking force were higher for Gag^M^ + gp140HA_2_tr and Gag^M^ + gp120HA_2_ than for Gag^M^ + gp150. The particle height was smaller for Gag^M^ + gp120HA_2_ than for Gag^M^ + gp150 and Gag^M^ + gp140HA_2_tr. Possible mechanisms underlying the increase of the particles’ stiffness and mechanical strength are the inclusion of the influenza virus HA transmembrane domain in the HIV Env protein, and the lower expression and packing density of Env in Gag^M^ + gp140HA_2_tr and Gag^M^ + gp120HA_2_ compared to Gag^M^ + gp150 found previously. Upon confirmation, the proposed mechanisms offer potential to tailor the mechanics of HIV virus-like particles and guide mechanical interactions between VLPs and host cells towards improving vaccines.

## 1 Introduction

The maturation of wild-type HIV involves the cleavage of the gp160 envelope protein into gp120 and gp140 which contains the cytoplasmic tail domain. For immature wild-type HIV, truncation of the cytoplasmic tail domain improves the fusion of the virion with the host cell (Murakami et al., 2004; Wyma et al., 2004) and reduces the virion’s stiffness (Kol et al., 2007). Pang et al. (2013) demonstrated that an increase in virion stiffness impaired viral entry activity, confirming the link between virion stiffness and entry ability and suggesting that virion stiffness may be an alternative target for the development of viral treatments.

Virus-like particles (VLPs) share some identical viral components with their wild-type counterparts but lack the infectious viral genomes. This absence allows VLPs to potentially replicate the entry process into host cells without providing the cells with the ability to produce infectious virions, thereby providing a safer alternative for immunocompromised individuals (Cervera et al., 2019; Nooraei et al., 2021). VLPs have been used for vaccines, drug delivery, gene therapy, and diagnostics (Fuenmayor et al., 2017; Nooraei et al., 2021; Rohovie et al., 2017; Zdanowicz and Chroboczek, 2016). For gene therapy and drug delivery, the selective entry of VLPs into specific cells can be induced by modifying the VLPs’ surface proteins (Nooraei et al., 2021). VLPs can be derived easily in laboratory settings using various cell types (Cervera et al., 2019; Fuenmayor et al., 2017; Nooraei et al., 2021) derived from mammals, plants (Greco et al., 2007; Vahdat et al., 2021), insects (Puente-Massaguer et al., 2021), and yeast (Sakuragi et al., 2002). VLPs have been used successfully as commercial vaccines for hepatitis B (Donaldson et al., 2018; Vahdat et al., 2021) and human papillomaviruses (Donaldson et al., 2018; Wang and Roden, 2013). VLPs offer an advantage over traditional vaccines due to their similar shape and size to virus particles, as well as their ability to elicit a similar immune response (Cervera et al., 2019; Gonelli et al., 2019; Nooraei et al., 2021). For HIV, VLPs present a promising vaccine modality as they present the envelope protein (Env) in a trimeric conformation, mimicking the wild-type virus (Gonelli et al., 2019).

Quantifying the VLP’s interactions with cells and their environment is essential to improve the understanding and utilisation of VLPs (Armanious et al., 2023). As it is for virions, the interaction with host cells is largely affected by the VLP’s mechanical properties and can influence the entry ability or genome release of the VLP (Buzon et al., 2020; Marchetti et al., 2016; Zandi et al., 2020). In cases where VLPs are used for the delivery of drugs or genetic material, their stability can influence disassembly and entry into host cells (Brunk and Twarock, 2021; Buzon et al., 2020; Marchetti et al., 2016). For applications where VLPs are used as nanoreactors, the mechanics of these VLPs influences their suitability and efficiency (Su et al., 2023). Furthermore, the capsid rigidity impacts the ability of the VLPs to withstand physical and chemical stresses during transport, storage and administration (Schumacher et al., 2018; Vishwakarma et al., 2024; Wubshet et al., 2024). Collett et al. (2019) reported significant differences in size amongst different hepatitis C VLP genotypes and a low elastic modulus of all VLP genotypes compared to other viruses and VLPs. They attributed these findings to the loosely packed capsid proteins typical of the hepatitis B virus. Armanious et al. (2023) showed that modification of VLPs for delivery to a mucosal environment through fusion of peptides and chemical conjugation of polymers induced considerable changes in the VLP’s biophysical properties. Thus, the mechanics of VLPs has an impact on their suitability for the desired application and studying the VLP mechanics can help improve future vaccine design (Radiom et al., 2023).

Replacing different sections of the stalk domain of Env (gp41) with corresponding components from other viral glycoproteins has been shown to enhance the density of Env spikes on both the cell membrane and the surface of virus-like particles (VLPs). Chapman et al. (2020) created chimaeric Env timers by substituting the transmembrane and cytoplasmic domains of HIV-1 Env with the corresponding regions from the influenza H5 hemagglutinin (HA) (gp140HA_2_tr), and by replacing the entire gp41 region of Env with the HA_2_ subunit of HA (gp120HA_2_). Rabbits inoculated with DNA and MVA expressing gp140HA_2_tr or gp150 induced autologous Tier 2 neutralising antibodies. However, those expressing the gp120HA_2_ Env did not. They then investigated the impact of including chimaeric trimers of influenza and HIV into HIV-1-like particles on Env spike density and antibody binding and demonstrated a difference in the binding of a monoclonal antibody to a V1/V2 quaternary epitope. This epitope was not detected in the vaccines expressing gp120HA_2_ Env. However, the effect of such a chimaera on the mechanical and morphological properties of HIV-like particles is unknown. Hence, the current study aimed to quantify the mechanical and morphological properties of three HIV-1 VLP types, which share the same Gag protein shell but differ in their trimeric glycoprotein envelopes, using atomic force microscopy, transmission electron microscopy, and finite element modelling.

## 2 Materials and methods

### 2.1 Virus-like particle (VLP) preparation

The three VLP types used in this study resemble HIV virions and have the same Gag protein shell and trimeric glycoprotein envelope as HIV, but they lack viral RNA. The plasmid DNA Gag^M^ used to produce the Gag protein shell contains an HIV-1 subtype C Gag mosaic gene insert that produces a protein that assembles efficiently in cultured cells (Chapman et al., 2017). The glycoprotein trimers are the viral ligands responsible for binding with the host cell, facilitating viral infection (Haywood, 1994). The furin cleavage site of gp160 was replaced with a flexible linker (FL) that is cleavage-independent to stabilise the linkage of gp120 to gp41 (Sharma et al., 2015). The cytoplasmic tail domain was truncated to gp150, promoting increased glycoprotein expression (Aldon et al., 2018). The gp150 trimer was combined with the Gag protein shell to produce the first VLP type Gag^M^ + gp150 (van Diepen et al., 2019). Two chimaeric VLPs were created by replacing all or part of the stalk domain of the envelope protein of HIV (Env) with sequences from influenza virus A H5 haemagglutinin (Chapman et al., 2020) (Figure 1). The entry-inducing domain of HIV (i.e., gp120) was retained in the VLPs to maintain the exact entry mechanism of HIV. The latter is essential because the human body will develop specific antibodies to the surface proteins of the viral particle, which should prevent future viral infections. Using the Gag protein shell and the chimeric trimer, the VLP types Gag^M^ + gp140HA_2_tr and Gag^M^ + gp120HA_2_ were produced.

**Figure 1.**
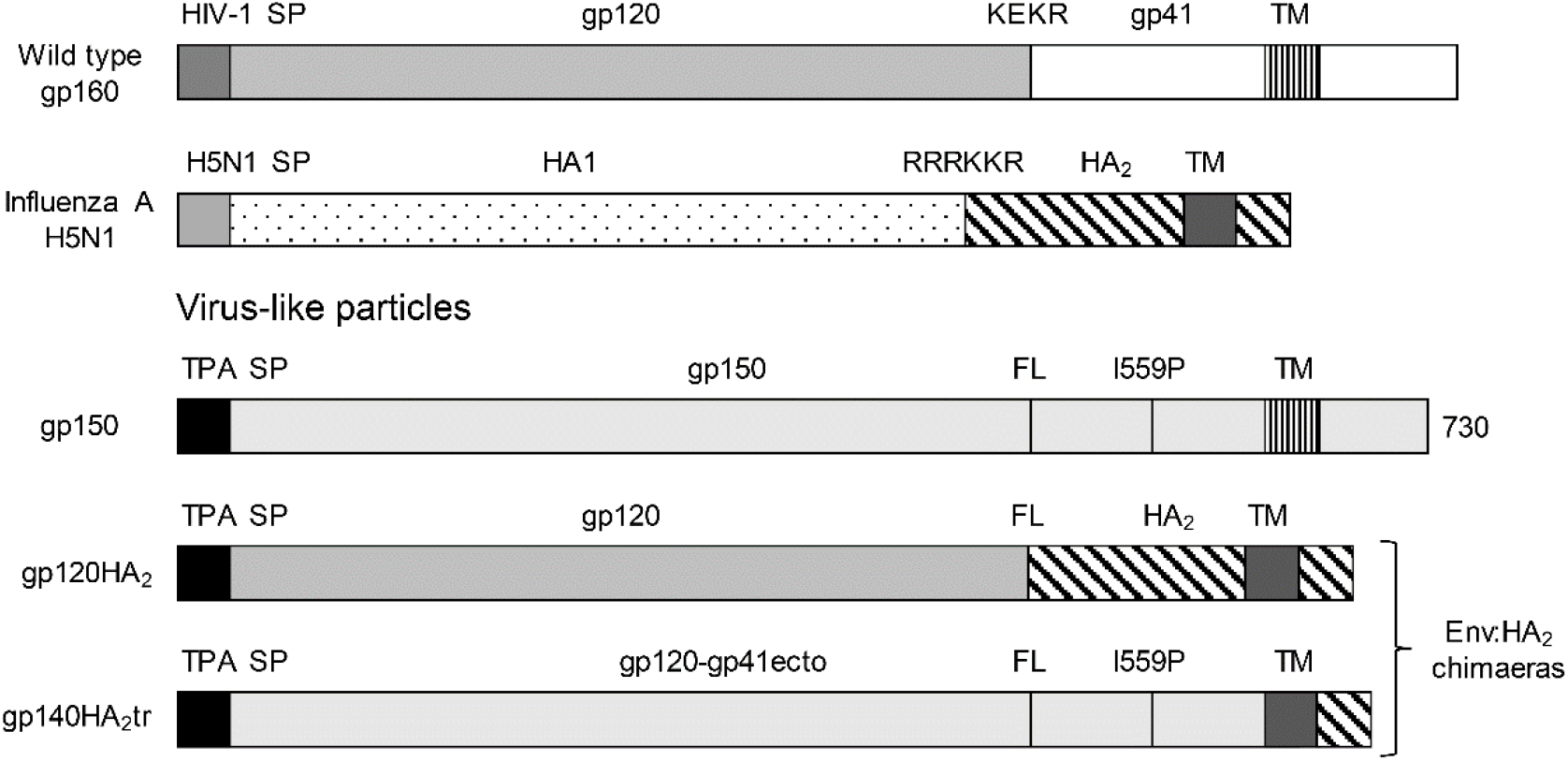
Schematic representations of wildtype HIV-1 gp160, influenza hemagglutinin (HA), gp150, gp120HA_2_, and gp140HA_2_tr. The gp160 sequence was truncated at amino acid residue 730 to generate gp150, the native signal peptide sequence (SP) was replaced with the human tissue plasminogen activator leader sequence (TPA), the furin cleavage motif, KEKR, was replaced with a flexible linker (FL), and an I559P mutation was introduced. Adjusted from (Chapman et al., 2020).

The VLP preparation included transfecting the host cells, isolating virus-like particles from the host cell cultures, and further purifying the particle suspension.

#### 2.1.1 Cell transfection

Human embryonic kidney cells (HEK293T, ATCC®, Manassas, USA, CRL-3216^TM^) were seeded in a T75 flask coated with poly-l-lysine (PLL, Sigma-Aldrich, Burlington, USA) and incubated at 37°C and 5% CO_2_. When the cells reached 70-80% confluency, the media was replaced with 12 ml Dulbecco’s modified Eagle’s medium (DMEM, Lonza, Basel, Switzerland) with 10% foetal calf serum (FCS, Gibco™, ThermoFisher Scientific, Waltham, USA), 5 ml Penicillin-Streptomycin (P/S, Gibco™, ThermoFisher Scientific) and 10 mM HEPES (4-(2-hydroxyethyl)-1-piperazineethanesulfonic acid, Gibco™, ThermoFisher Scientific) with pH 7.4. A transfection mixture was prepared by adding 30 µg of the appropriate plasmid DNA (15 µg Env or EnvHA_2_ chimaeras + 15 µg Gag^M^) to 750 µl of DMEM, adding 90 µl of polyethylenenimine (PEI, Sigma-Aldrich) to 750 µl DMEM and then adding the PEI mixture to the DNA mixture and incubating at room temperature for 20 minutes. The transfection mixture was added to the HEK293T cells and incubated for three days. During the three-day incubation, a total of 8 ml of DMEM + 10% FCS + P/S + 10 mM HEPES pH 7.4 was added to the flask.

#### 2.1.2 Particle isolation

To isolate the particles from the transfected HEK293T cells, the supernatant was collected and spun down in a 50 ml centrifuge tube (Corning®) at 2500 g and 4°C for 5 minutes. The supernatant was collected, filtered through a 0.2 µm syringe filter (Corning®, New York, USA), and placed in a 50 ml Oak Ridge centrifuge tube (Nalgene®, Rochester, USA). A 5 ml 12% OptiPrep (60%, Sigma-Aldrich) cushion solution was underlaid at the bottom of the 50 ml Oak Ridge centrifuge tube, and the tube was spun at 50228 g and 4°C for 45 minutes. The supernatant was then discarded, and the pellet was suspended in 100 µl tris-buffered saline (TBS, Gibco™, ThermoFisher Scientific) for TEM samples or phosphate-buffered saline (PBS, Gibco™, ThermoFisher Scientific) for AFM samples and stored at -80°C.

#### 2.1.3 Particle suspension cleaning

The particle solution was cleaned using a PD Minitrap™ G-25 column (GE Healthcare, Chicago, USA) with a gravity protocol, and intervals of 40 µl fractions were collected. Fractions 4 and 5 were typically used for experimentation as they exhibited the least background debris.

### 2.2 Validation of VLP production using Western blots

15 µl of VLP sample and 5 µl of 4x protein loading dye were prepared and boiled at 95°C for 5 minutes. 5 µl of molecular weight marker and 20 µl sample were loaded into an SDS PAGE tank (BioRad MiniPROTEAN® Tetra Vertical Electrophoresis system, Hercules, USA) with a 10% resolving gel. The system was run at 200 V until the sample had run through the gel. The gel was removed and submerged in PBS with 0.1% Tween® 20 (PBST, Sigma-Aldrich). The gel was transferred using the iBlot™ 2 Gel Transfer Device (ThermoFisher Scientific), removed, and blocked in 5% non-fat milk (Sigma-Aldrich) in PBS (blocking buffer) solution for 20 minutes on an orbital shaker. The blocking buffer was removed, and the membrane was incubated overnight on an orbital shaker in a solution containing the primary antibodies, Goat anti-HIV-1 gp120 (Bio-Rad; 5000-0557) and goat anti-HIV-1 p24 (Gag) (Bio-Rad; 4999-9007), diluted in the blocking buffer. The primary antibody solution was removed, and the membrane was washed thrice with PBST on the orbital shaker for 5 minutes. 2 µl of mouse Monoclonal Anti-Goat/Sheep IgG Alkaline Phosphatase antibody produced in mouse (Sigma-Aldrich; A8062) was diluted in 20 ml of blocking buffer and incubated at room temperature for a minimum of 1 hour with shaking. The secondary antibody solution was removed, and the membrane was washed thrice with PBST on the orbital shaker for 5 minutes. The PBST was removed, 5 ml of KPL BCIP/NBT phosphatase substrate (SeraCare, Milford, USA) was pipetted directly onto the membrane, and the membrane was left in the dark until bands were detected. The reaction was stopped by submerging the membrane in PBST and air-drying overnight in the dark.

### 2.3 Transmission electron microscopy (TEM) imaging and morphometric assessment of VLP

s TEM imaging was performed to determine the outer diameter of the VLP and the wall thickness of the Gag protein shell.

A carbon grid was floated on a droplet of the sample for 30 seconds and then blotted on tissue paper. The grid was washed with 2% uranyl acetate, blotted with tissue paper twice, incubated on a droplet of 2% uranyl acetate for 1 minute, blotted with tissue paper again, and air-dried for 5–10 minutes. VLP samples were imaged using an FEI Tecnai F20 electron microscope (Hillsboro, USA) operated at 200 kV under low-dose conditions. Images were captured on a 4,096 x 4,096 pixel CCD camera at x100,000 magnification (Manole et al., 2012; Patel, 2015).

The outer diameter and wall thickness were measured for three particles of each VLP type using Fiji (Schindelin et al. 2012). The mean wall thickness was used to calculate the elastic modulus (see Section 2.4.2), and the mean particle diameter and wall thickness values were used for the finite element (FE) analysis (see Section 2.5).

### 2.4 AFM assessment of VLPs

Glass coverslips were used as substrates. Clean coverslips were submerged in a solution of 25 g potassium chloride, 25 ml Milli-Q water, and 200 ml 96% ethanol for 15 hours in a sealed glass jar. The coverslips and jar were then rinsed with Milli-Q, partially dried using argon, and dried completely overnight in the glass jar. 1 ml of hexamethyldisilazane (HMDS) was added to the bottom of the glass jar, sealed and incubated with the glass coverslips for 15 hours to generate a hydrophobic surface on the glass coverslips. Thereafter, the HMDS was allowed to evaporate in a fume hood for 3 hours (Roos, 2011; Snijder et al., 2012).

A glass coverslip was placed on the AFM stage, and 10 µl of the sample solution was applied to the coverslip and incubated at room temperature for 15 minutes. The AFM stage was then placed in the AFM, and 100 µl of PBS was added to the sample well.

All AFM experiments were conducted with a JPK NanoWizard® 3 Ultra Speed AFM (JPK Instruments AG, Berlin, Germany) and cantilever qp-BioAC CB3 (Uniqprobe®, Nanosensors™, Neuchatel, Switzerland) with a manufacturer-specified spring constant of 0.03 - 0.09 N/m and a tip radius smaller than 10 nm. The specific spring constant of each cantilever tip was measured at the start of each experiment using the calibration tool on the JPK NanoWizard® software, and the average cantilever stiffness was 0.072 ± 0.004 N/m.

A sample included (i) force spectroscopy imaging of the entire sample to identify individual particles, (ii) nanoindentation of individual particles, and (iii) repeated force spectroscopy imaging of each identified particle to capture possible morphological changes of the particles during nanoindentation. 75 particles were assessed: n = 19 for Gag^M^ + gp150, n = 31 for Gag^M^ + gp140HA_2_tr, and n = 25 for Gag^M^ + gp120HA_2_.

All raw AFM data were exported from JPKSPM Data Processing 6.1.79 (JPK Instruments AG) and imported into MATLAB® R2021a for analysis.

#### 2.4.1 Morphological assessment using force spectroscopy mode

The particle height was determined by the maximum pixel intensity after applying the plane fitting function in JPKSPM Data Processing 6.1.79 (JPK Instruments AG). Images were cropped before the height measurement to eliminate the effects of background debris, imaging noise, or multiple particles, ensuring the correct height was measured.

#### 2.4.2 Mechanical assessment using nanoindentation

The experiment involved the nanoindentation of a VLP with a peak force setpoint of 5 nN. The spring constant of the VLP was determined from linear fits to two regions of the force-displacement curve, i.e. k_R1_ for the region defined by a force between 0.1 nN and 0.2 nN, and k_R2_ for the force region between 0.2 nN and the breaking point (Li et al., 2011; Schaap et al., 2012). The breaking force was defined as a decrease in the force value for two consecutive data points, accompanied by an increase in displacement.

The force-displacement curve represents the system’s total spring constant, k_t_. The sample spring constant, k_s_, is calculated using Hooke’s law of two springs with the known cantilever spring constant, k_c_:

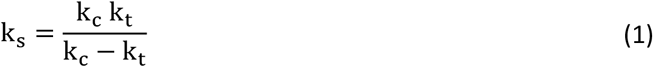

The thin shell model was used to determine the elastic modulus E of the VLPs (Michel et al., 2006; Roos, 2011; Roos et al., 2009):

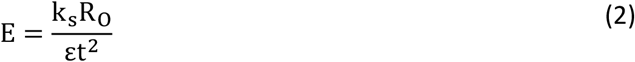

where k is the spring constant, R_O_ is the VLP outer radius, *ε* is a proportionality factor, and t is the wall thickness of the VLP. The wall thickness was obtained morphometrically from TEM images of the three VLP types as described above. The proportionality factor was approximated as 1 (Gibbons and Klug, 2007; Michel et al., 2006; Roos, 2011).

#### 2.4.3 Assessment of particle movement during nanoindentation

The movement of a particle during nanoindentation can affect the force-displacement data and lead to misinterpretation, e.g. as particle failure. To check for any movement of a particle on the substrate, the AFM images of the particle before and after indentation were imported into Fiji (Schindelin et al., 2012), aligned using the DS4H Image Alignment plugin (Bulgarelli et al., 2019) with three background debris items, and visually inspected. This procedure confirmed the absence of particle movement for all nanoindentation experiments (Figure S1).

### 2.5 Finite element simulation of VLP nanoindentation

#### 2.5.1 Geometry

The VLP was modelled in Abaqus/CAE 2022 (Simulia, Dassault Systèmes) and represented by a spherical shell geometry with outer and inner diameters obtained from morphometric analysis of TEM images (Section 2.3), i.e. the outer diameter of 172 nm for Gag^M^ + gp150, 179 nm for Gag^M^ + gp140HA_2_tr, and 169 nm for Gag^M^ + gp120HA_2_. and inner diameter of 138 nm for Gag^M^ + gp150, 144 nm for Gag^M^ + gp140HA_2_tr, and 130 nm for Gag^M^ + gp120HA_2_.

The cantilever tip and substrate were geometrically modelled as a solid sphere with a radius of 10 nm and a rectangular cuboid with dimensions of 200 x 200 x 20 nm, respectively (Abaqus CAE 2022, Simulia, Dassault Systèmes).

#### 2.5.2 Mesh specifications

The VLP was meshed using the sweep technique, with the sweep path around the sphere’s circumference, refining the mesh from the mid-circumference towards the contact region with the indenter. A hex-dominated mesh with hexahedral elements (C3D8I) and linear wedge elements (C3D6) was used. The three VLP types had slightly different element numbers of 278,976 for Gag^M^ + gp150, 282,692 for Gag^M^ + gp140HA_2_tr, and 278,786 for Gag^M^ + gp120HA_2_, due to the different dimensions of the three VLP types.

The cantilever tip and substrate were meshed with 2,208 and 16, respectively, 8-node linear brick elements with incompatible modes (C3D8I).

A mesh convergence study was conducted to determine an optimal mesh size. Convergence was reached at approximately 165,000 elements, with a difference in the force value of less than 5% for all simulations with a finer mesh (i.e. more than 165,000 elements), see Table S1.

#### 2.5.3 Material models

The VLP was modelled as an isotropic linear-elastic material with a Poisson ratio of 0.4 (Ahadi et al., 2013). The value of the elastic modulus was selected such that the numerically predicted behaviour of the VLP resembled the experimental behaviour (see Section 2.5.5).

#### 2.5.4 Boundary conditions, contact, and loading

Rigid body constraints were applied to the indenter sphere and substrate to prevent the deformation of these parts. Sliding, frictionless, and normal hard contact was implemented between the virion and the indenter, as well as between the virion and the substrate. Further boundary conditions constrained the substrate fully, whereas the virion was allowed to move in the indentation direction. A displacement boundary condition in the z-direction assigned to the indenter sphere represented the displacement of the indenter tip during indentation. The maximum displacement, d_ind_, was determined for each VLP type with the experimentally determined spring constant of the VLP, k_R1_, and a prescribed maximum indentation force of F_ind_ = 0.2 nN, i.e. d_ind_ = F_ind_/k_R1_, which was 16.60 nm for Gag^M^ + gp150, 8.90 nm for Gag^M^ + gp140HA_2_tr and 7.56 nm for Gag^M^ + gp120HA_2_.

#### 2.5.5 Simulations and data recording

Simulations of the nanoindentation process were performed repeatedly, adjusting the value of the elastic modulus of the VLP from an initial value obtained from the thin shell model (Section 2.4.2), until the indentation force through the centre of the indenter sphere was F_ind_ = 0.2 ± 0.0025 nN at the maximum indentation displacement, d_ind_.

### 2.6 Statistical analysis

All statistical analyses were performed using IBM SPSS Statistics® 25. Generalised estimating equations were used to determine differences between the VLP types. Spearman’s Rank-Order Correlation was used for correlation tests. Statistical significance was assumed to be approached for p ≤ 0.05. All error values are given as a standard error of the mean (SEM).

## 3 Results

### 3.1 Validation of VLP production using western blots

The western blots show positive staining of the Gag and Env proteins for the three VLP types (Figure 2a). For VLPs produced using only Gag^M^, there was positive staining of only Gag proteins; for samples transfected with gp150 only, there was positive staining of the Env proteins. No staining was apparent for the non-transfected control group.

**Figure 2.**
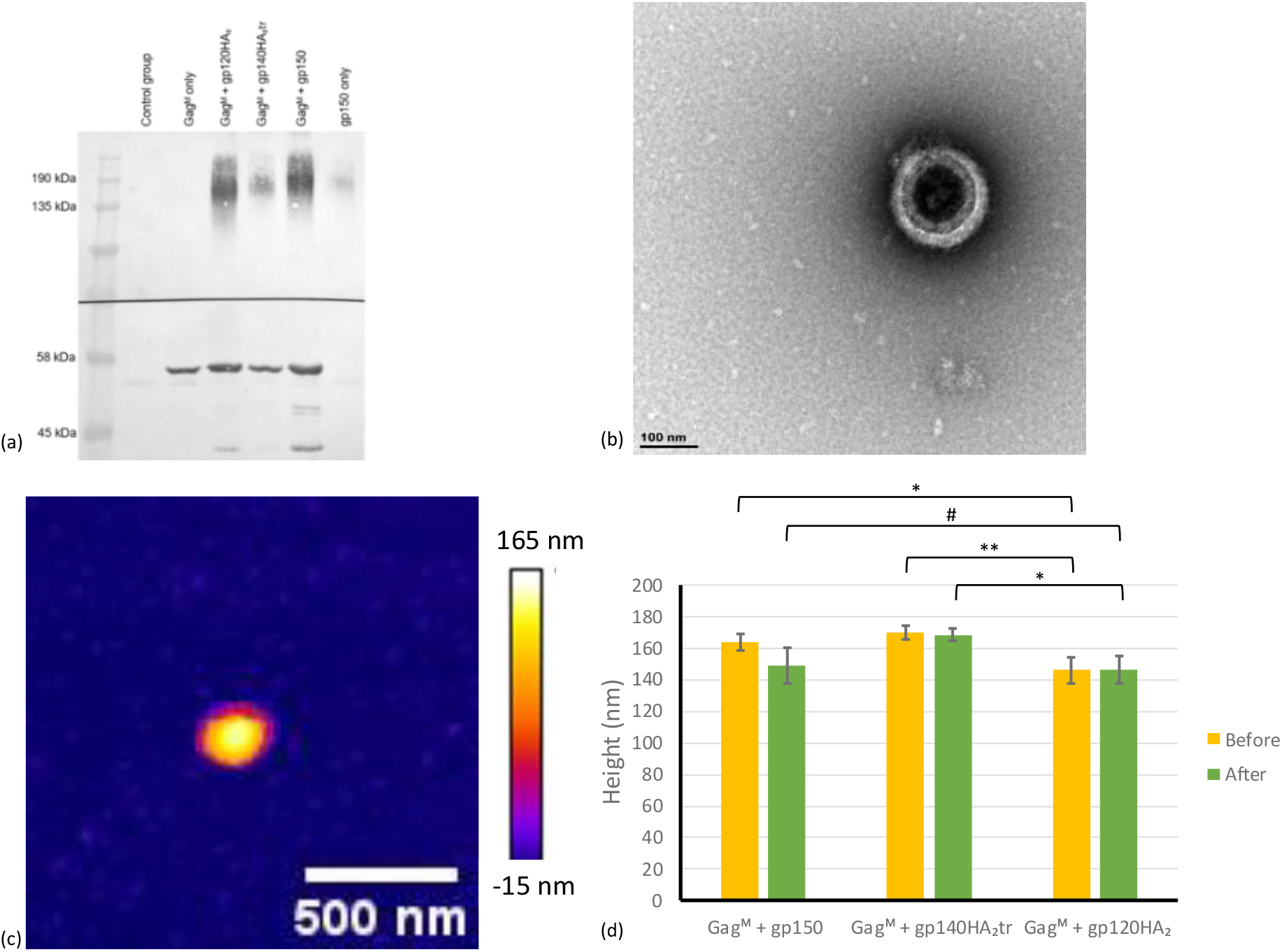
(a) Western blot of the VLPs collected after spinning in the ultracentrifuge shows positive staining of the Gag and Env proteins for the three VLP types. (b) Transmission electron micrograph and (c) atomic force micrograph of a VLP sample after cleaning with the column (4^th^ fraction) show successful isolation of the VLPs. (d) Mean height of the VLP types determined with AFM before and after nanoindentation. [n = 19 for Gag^M^ + gp150, n = 31 for Gag^M^ + gp140HA_2_tr, and n = 25 for Gag^M^ + gp120HA_2_. # p = .05001, * p < .05, ** p < .01. Error bars indicate the standard error of the mean (SEM)].

### 3.2 Morphological properties of VLPs

Imaging the samples using TEM and AFM confirmed that spherical particles with a diameter in the 100 nm range were produced (Figure 2 b & c). The AFM assessment indicated a VLP height before and after nanoindentation of 164 ± 5 nm and 149 ± 11 nm (n = 19, p = .826) for Gag^M^ + gp150, 170 ± 4 nm and 168 ± 4 nm (n = 31, p = .332) for Gag^M^ + gp140HA_2_tr, and 146 ± 9 nm and 146 ± 9 nm (n = 25, p = .739) for Gag^M^ + gp120HA_2_ (Figure 2 d). There was a significant height difference between Gag^M^ + gp120HA_2_ and the other two particle types.

Morphometric TEM image analysis provided an outer diameter, inner diameter and wall thickness of 172 ± 10 nm, 138 ± 11 nm, and 17.3 ± 1.5 nm for Gag^M^ + gp150 (n = 3), 179 ± 3 nm, 144 ± 2 nm and 17.7 ± 0.3 nm for Gag^M^ + gp140HA_2_tr (n = 3), and 169 ± 4 nm, 130 ± 5 nm and 19.7 ± 1.3 nm for Gag^M^ + gp120HA_2_ (n = 3). The outer diameter values based on TEM were not significantly different from the VLP height measured with AFM before nanoindentation (p = .264 for Gag^M^ + gp150, p = .732 for Gag^M^ + gp140HA_2_tr, and p = .060 for Gag^M^ + gp120HA_2_).

### 3.3 Mechanical properties of VLPs

The results of the mechanical assessment include the spring constant, the breaking force, and the elastic modulus.

#### 3.3.1 Spring constant

The spring constants of the VLP for the two regions of the force-displacement curve (Figure S2) are k_R1_ = 0.012 ± 0.002 N/m and k_R2_ = 0.014 ± 0.002 N/m for Gag^M^ + gp150, k_R1_ = 0.022 ± 0.004 N/m and k_R2_ = 0.08 ± 0.02 N/m for Gag^M^ + gp140HA_2_tr, and k_R1_ = 0.026 ± 0.005 N/m and k_R2_ = 0.14 ± 0.05 N/m for Gag^M^ + gp120HA_2_ (Figure 3 a).

**Figure 3.**
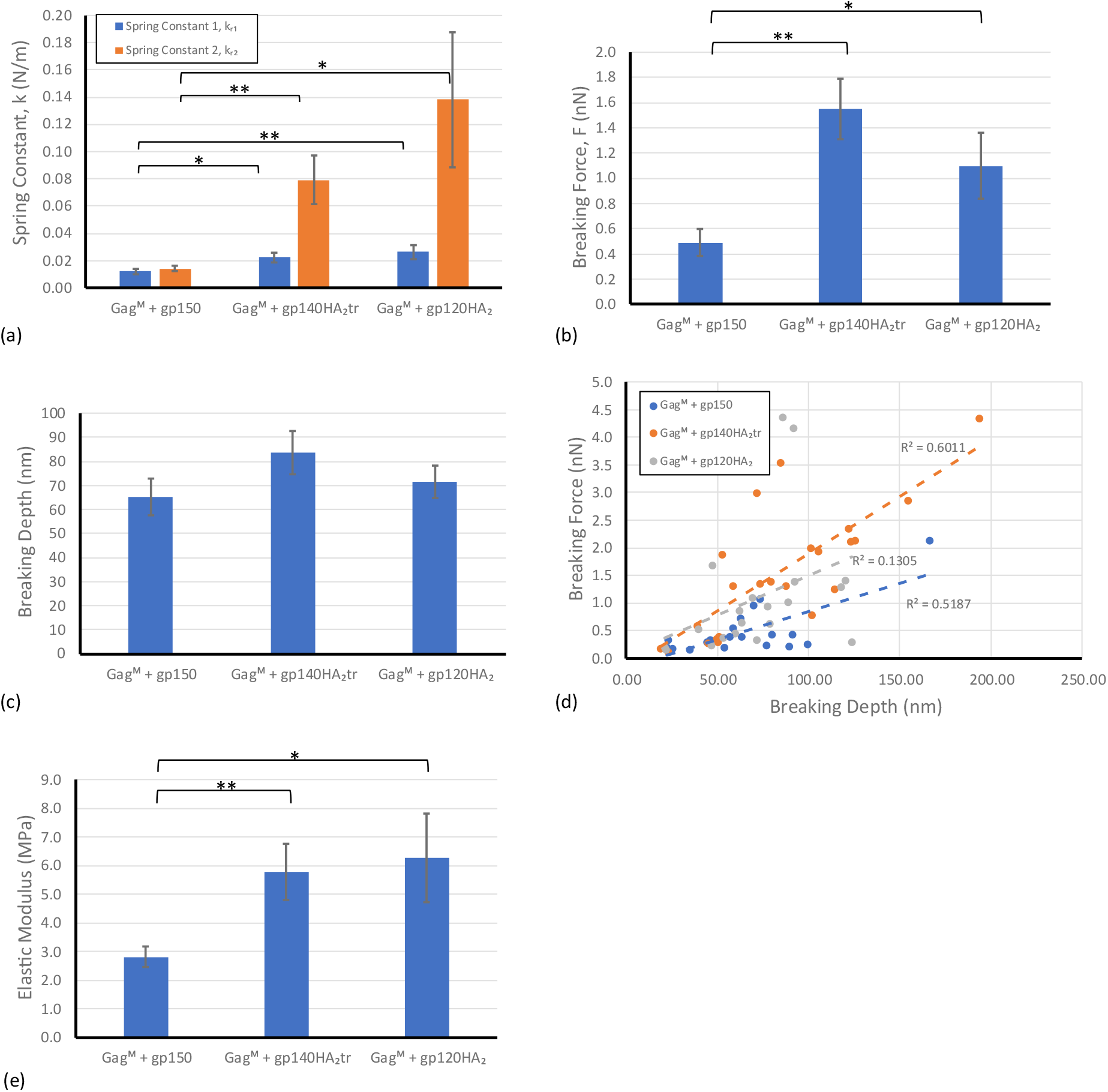
(a) The spring constant of the three VLP types for the two different slope fits (indicated with the slope number above each bar). (b) The mean breaking force for the three different VLP types. (c) The mean breaking depth for the three different VLP types. (d) A scatter plot of breaking force versus breaking point depth for each virion with a linear trendline for all three VLP types. (e) The elastic modulus of the three VLP types determined using the thin shell model. (*p < .05, **p < .01. Error bars indicate SEM.)

A significant difference between Gag^M^ + gp150 compared to the other two VLP types for k_R1_ (p = .010 compared to Gag^M^ + gp140HA_2_tr, p = .004 compared to Gag^M^ + gp120HA_2_ and k_R2_ (p < .001 compared to Gag^M^ + gp140HA_2_tr, p = .011 compared to Gag^M^ + gp120HA_2_ . The difference in the spring constant of Gag^M^ + gp140HA_2_tr and Gag^M^ + gp120HA_2_ was not significant (p = .559 for k_R1_ and p = .274 for k_R2_).

#### 3.3.2 Breaking force and depth

The mean breaking force was F_b_ = 0.5 ± 0.1 nN for Gag^M^ + gp150, F_b_ = 1.6 ± 0.2 nN for Gag^M^ + gp140HA_2_tr, and F_b_ = 1.1 ± 0.3 nN for Gag^M^ + gp120HA_2_ is (Figure 3 b). The breaking force of Gag^M^ + gp150 was significantly lower than that of Gag^M^ + gp140HA_2_tr (p < .001) and Gag^M^ + gp120HA_2_ (p = .030). The proportion of broken VLPs varies with VLP type, with breakage of 100% (19/19) for Gag^M^ + gp150, 74% (23/31) for Gag^M^ + gp140HA_2_tr and 80% (20/25) for Gag^M^ + gp120HA_2_.

The depth at which the VLPs broke was d_b_ = 65.1 ± 7.7 nm for Gag^M^ + gp150, 83.8 ± 9.7 nm for Gag^M^ + gp140HA_2_tr, and 71.6 ± 6.6 nm for Gag^M^ + gp120HA_2_ (Figure 3c). The differences in breaking depth between the VLP types were not significant (p > .1).

The correlation between the breaking force and breaking depth was strong for Gag^M^ + gp150 (r = .720, p = .024) and Gag^M^ + gp140HA_2_tr (r = . 5, p < .001, but weak for Gag^M^ + gp120HA_2_ (r = .361, p = .011) (Figure 3 d).

#### 3.3.3 Elastic modulus based on the thin shell model

The wall thickness of t = 18.2 nm (average for the three VLP types from TEM images) was used to determine the elastic modulus using the thin shell model, Eq. (2), with k_R1_ for each VLP type. The elastic modulus was E = 2.8 ± 0.4 MPa for Gag^M^ + gp150, E = 5.8 ± 1.0 MPa for Gag^M^ + gp140HA_2_tr, and E = 6.3 ± 1.55 MPa for Gag^M^ + gp120HA_2_ (Figure 3 e). The elastic modulus of Gag^M^ + gp150 was significantly lower than those of Gag^M^ + gp140HA_2_tr (p = .004) and Gag^M^ + gp120HA_2_ (p = .027), whereas there was no significant difference between Gag^M^ + gp140HA_2_tr and Gag^M^ + gp120HA_2_ (p = .791).

#### 3.3.4 FE-based elastic modulus

The FE simulations with the optimised elastic modulus yielded a satisfactory replication of the *in vitro* experiments, using displacement control with a similar indentation depth d_ind_ corresponding to the indentation force of F_ind_ = 0.2 nN (Figure 4).

**Figure 4.**
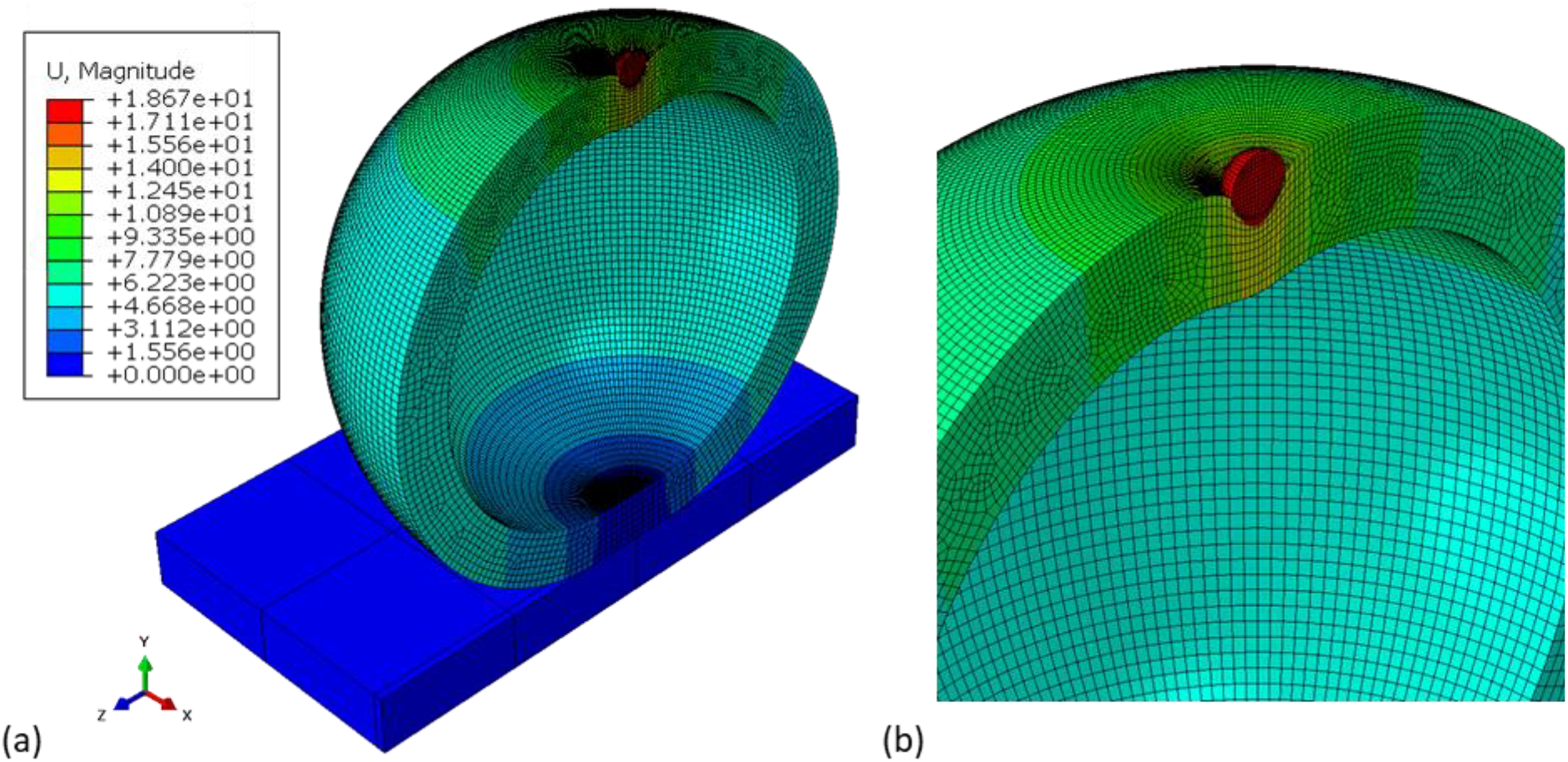
(a) Symmetric cross-sectional view of the FE model of indenter sphere, VLP, and substrate illustrating the deformed Gag^M^ + gp150 VLP geometry and contours of displacement at an indentation depth corresponding to an indentation force of F_ind_ = 0.2 nN. (The displacement is given in nm). (b) A close-up view of the contact region of the rigid indenter and VLP.

**Figure 5.** FE-predicted indentation force versus displacement for the optimised value of the elastic modulus for each VLP type up to an indentation force of F_ind_ = 0.2 nN.

The FE simulations predicted an elastic modulus of E = 1.9 MPa for Gag^M^ + gp150, E = 4.5 MPa for Gag^M^ + gp140HA_2_tr, and E = 5.6 MPa for Gag^M^ + gp120HA_2_, which was slightly lower than the respective values obtained from the thin-shell model.

### 3.4 Correlation of spring constant and height of VLPs

There was no consistent correlation between the spring constant k_R1_ and height for the three VLP types. For Gag^M^ + gp150, there was a moderate negative correlation (r = -.655, p = .002), whereas there was a weak and statistically not significant positive correlation for Gag^M^ + gp140HA_2_tr (r = .071, p = .704) and Gag^M^ + gp120HA_2_ (r = .293, p = .155).

## 4 Discussion

The current study employed atomic force microscopy, transmission electron microscopy, and finite element analysis to investigate the mechanical properties of HIV-like particles and their variation with modifications to the Env membrane proteins.

The mechanical properties obtained suggest that modification of the Env protein by including the transmembrane or larger domain of influenza virus A HA can improve the mechanical stability of the VLPs, as both Gag^M^ + gp140HA_2_tr and Gag^M^ + gp120HA_2_ have a significantly higher spring constant and breaking force than Gag^M^ + gp150 (Table 1). The correlation between stiffness and virion height also indicated that Gag^M^ + gp140HA_2_tr and Gag^M^ + gp120HA_2_ have similar mechanical behaviour.

**Table 1.** Mechanical properties of VLPs.

|  | Spring constant (N/m) |  | Elastic modulus | Breaking force |
| --- | --- | --- | --- | --- |
| | $k_{R1}$ | $k_{R2}$ | (MPa)<sup>1</sup> | (nN) |
| Gag<sup>M</sup> + gp150 | $0.012 \pm 0.002$ | $0.014 \pm 0.002$ | 1.9 | $0.5 \pm 0.1$ |
| Gag<sup>M</sup> + gp140HA<sub>2</sub>tr | $0.022 \pm 0.004$ | $0.08 \pm 0.02$ | 4.5 | $1.6 \pm 0.2$ |
| Gag<sup>M</sup> + gp120HA<sub>2</sub> | $0.026 \pm 0.005$ | $0.14 \pm 0.05$ | 5.6 | $1.1 \pm 0.3$ |

<sup>1</sup> Elastic modulus from FE model.

Previous studies indicated that Env plays a role in regulating the stiffness of particles (Kol et al., 2007; Pang et al., 2013). Chapman et al. (2020) found a difference in the Env:Gag ratio for the three VLP types investigated in the current study, where Gag^M^ + gp150 VLP has the highest Env:Gag ratio and Gag^M^ + gp120HA_2_ has the lowest Env:Gag ratio. This difference, although not statistically significant, may indicate a difference in the expression or packing density of Env that inversely affects the stiffness of the particles observed in the current study. Currently, there is no data available on the stiffness of VLPs composed solely of the Gag protein, without a viral ligand, that could provide insight into how the Env proteins affect the stability of these VLPs.

Mature and immature HIV pseudovirions were reported to have stiffnesses of 0.22 N/m and 3.16 N/m, respectively, and elastic moduli of 440 MPa and 930 MPa (Kol et al., 2007). These values differ from those in the current study, which reported lower stiffness and elastic modulus. Genetic material contributes to the mechanics of virus particles (Carrasco et al., 2008; Michel et al., 2006). Hence, the differences in the genetic material of the VLPs in the current study and the pseudovirions used by Kol et al. (2007) may contribute to the difference in the mechanical properties. Other studies of VLPs, although of entirely different types, yielded similar results to the current work, such as an elastic modulus between 1.5 and 11.6 MPa for Hepatitis C VLPs (Collett et al., 2019) and between 6.2 and 21.3 MPa for SARS-CoV2 and dengue VLPs (Collett et al., 2023).

The thin shell model has been used previously to estimate the elastic modulus of spherical virions with a shell thickness of less than 10% of the virion diameter (Ivanovska et al., 2004; Roos et al., 2009). For small virion deformations, the thin shell model has also been found accurate for virions with thicker shells (Michel et al., 2006). However, as the VLPs in the current study have a shell thickness of approximately 20% of the VLP diameter, a finite element model is considered more robust and reliable than the thin shell model to determine the VLP elastic modulus. The finite element models yielded a slightly lower elastic modulus than the thin shell model for each VLP type.

AFM assessment determined a virion height before nanoindentation of 164 nm Gag^M^ + gp150, 170 nm for Gag^M^ + gp140HA_2_tr, and 146 nm for Gag^M^ + gp120HA_2_. After nanoindentation, the heights were 149 nm, 168 nm, and 146 nm for the three respective particles. No significant difference in particle size was observed before and after nanoindentation. The size of HIV virions has been reported to be 110 - 146 nm (Gentile et al., 1994) and 145 ± 15 nm (Briggs et al., 2003). HIV has also been reported to exhibit considerable variation in particle size, with individual virions reaching up to 200 nm (Briggs et al., 2003) and 240 nm (Kuznetsov et al., 2003). Although the average VLP sizes in the current study are larger than that of HIV, the particle height of the VLP types falls within the size range previously reported for HIV and is similar to a diameter of 172 ± 10 nm reported for HIV-like particles (Endress et al., 2008). The larger average VLP size may result from differences in the assembly processes of VLPs and HIV or from swelling of VLPs due to their internal osmotic pressure.

Despite apparent breaking points in the nanoindentation force-depth data, permanent fracture or buckling damage of the VLPs was not observed visually after nanoindentation. Image analysis also confirmed the absence of movement or slipping of VLPs during nanoindentation. A second nanoindentation procedure on a limited number of particles indicated differences in the force-depth curve and the particle stiffness compared to the first nanoindentation. This stiffness difference suggests that the particle undergoes permanent damage during the first indentation. Such damage may be caused by buckling and unbuckling of the particle due to the osmolarity difference between the fluid in the particle and the buffering media. The different osmolarities may cause the particle to fill up with fluid and regain its spherical shape instantaneously. A similar structural recovery after repeated nanoindentation was observed for SARS-CoV-2 virions by Kiss et al. (2021) and for T7 bacteriophages by Voros et al. (2017). Lastly, the breaking point in the force-depth curve may represent the puncturing of the virion shell with the cantilever probe (Nasto et al., 2013; Streltsov et al., 2023; Zhang et al., 2010). Further experiments and a larger sample size are needed to confirm and elucidate this phenomenon.

## 5 Conclusions

The current study determined the mechanical and morphological properties of three different HIV-VLPs. The results showed that including the influenza protein HA_2_ in the envelope structure of the HIV-like particles increased the breaking force and spring constant of the particles. This could indicate that including the influenza envelope proteins affects the mechanical behaviour of the VLPs and improves their structural integrity. These results demonstrate how mechanical factors can be utilised for designing future VLPs. Further work is required to quantify the expression and packing density of the envelope proteins and to determine how these parameters may be affected by the inclusion of the HA_2_ protein segments. Assessment of the Gag protein without Env will provide insight into the impact of the Env proteins on the mechanical properties of VLPs.

## Data Availability

Data and software code supporting this study are available on the University of ape Town’s institutional data repository ZivaHub under the DOI https://doi.org/10.25375/uct.28016702 as Kruse E, van Diepen M, Chapman R, Horn E, Abdalrahman T, Williamson A-L, Rybicki E, Roos WH, Franz T. Data for “Impact of modification of envelope proteins on the mechanical properties of HIV virus-like particles”. ZivaHub, 2025, DOI 10.25375/uct.28016702.

## Funding

The work was supported by the National Research Foundation of South Africa (grants UID92531 and UID93542 to TF and an Innovation Doctoral Scholarship to EK), the South African Medical Research Council (grant SIR328148 to TF), and the University of Cape Town (Doctoral Research Scholarship, KW Johnston Bequest Scholarship, and Murray-Jelks Scholarship to EK). This work was also supported by the South African Medical Research Council, with funds received from the South African Department of Science and Technology. ALW was funded by the National Research Foundation of South Africa (grant 64815).

## Competing Interests

The authors declare that they have no competing interests.

## Credit Author Contributions

E Kruse: Conceptualization, Data curation, Formal analysis, Investigation, Methodology, Project administration, Software, Validation, Visualization, Writing – Original Draft Preparation, Writing – Review & Editing

M van Diepen: Methodology, Supervision, Writing – Review & Editing

R Chapman: Methodology, Writing – Review & Editing

E Horn: Methodology, Writing – Review & Editing

T Abdalrahman: Methodology, Software, Writing – Review & Editing

A-L Williamson: Resources, Writing – Review & Editing

E Rybicki: Resources, Writing – Review & Editing

WH Roos: Investigation, Resources, Supervision, Writing – Review & Editing

T Franz: Conceptualization, Data curation, Funding acquisition, Methodology, Project administration, Resources, Supervision, Validation, Writing – Original Draft Preparation, Writing – Review & Editing

## Notes

https://doi.org/10.25375/uct.28016702

### Competing Interest Statement

The authors have declared no competing interest.

