## Supplemental material for "Impact of modification of envelope proteins on the mechanical properties of HIV virus-like particles"

Supplemental information

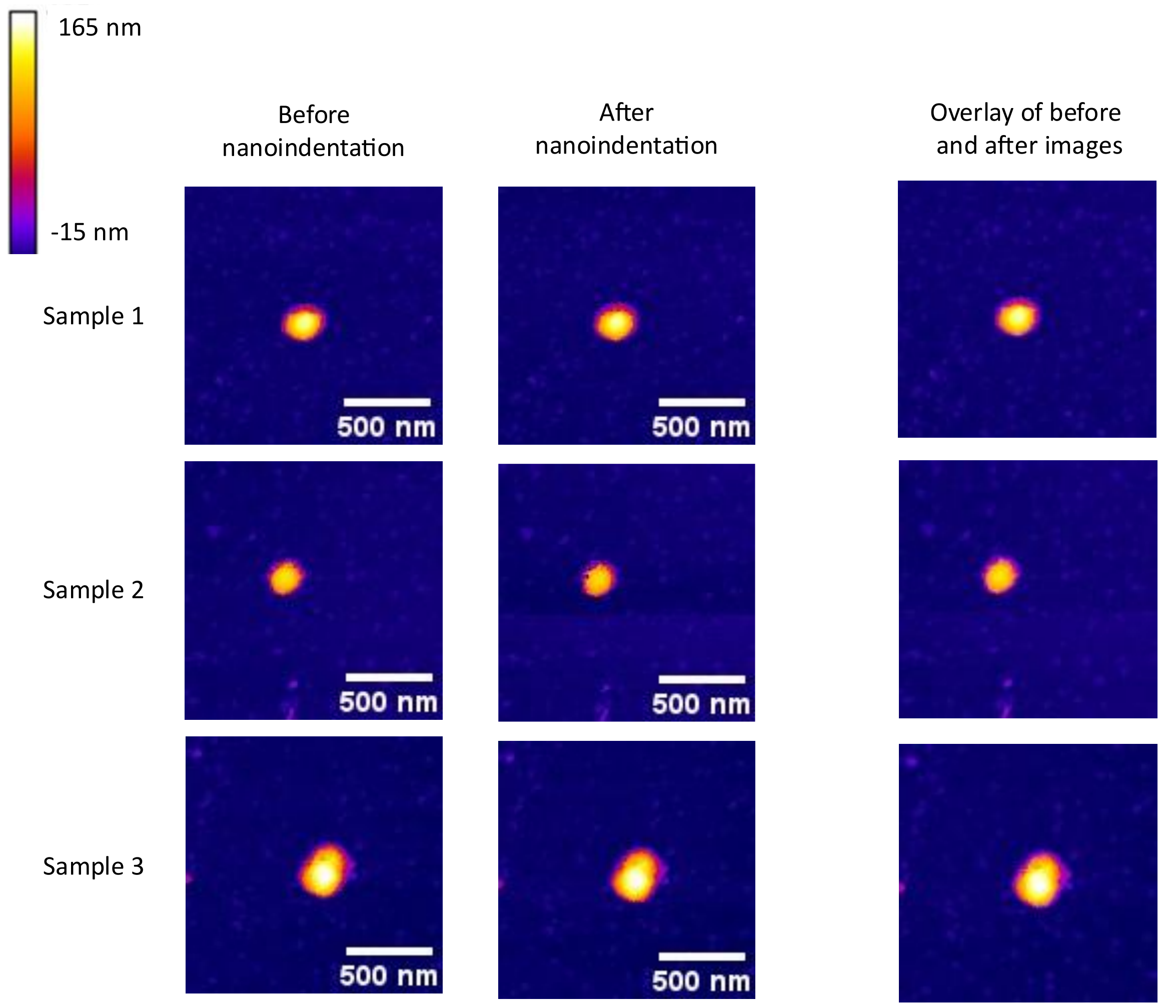


Figure S1. The images of three samples before and after nanoindentation as well as the two images aligned and overlayed. The height colour scale bar is the same for all the before and after nanoindentation images. The overlay images do not have a height scale as they are a composite of the before and after images.

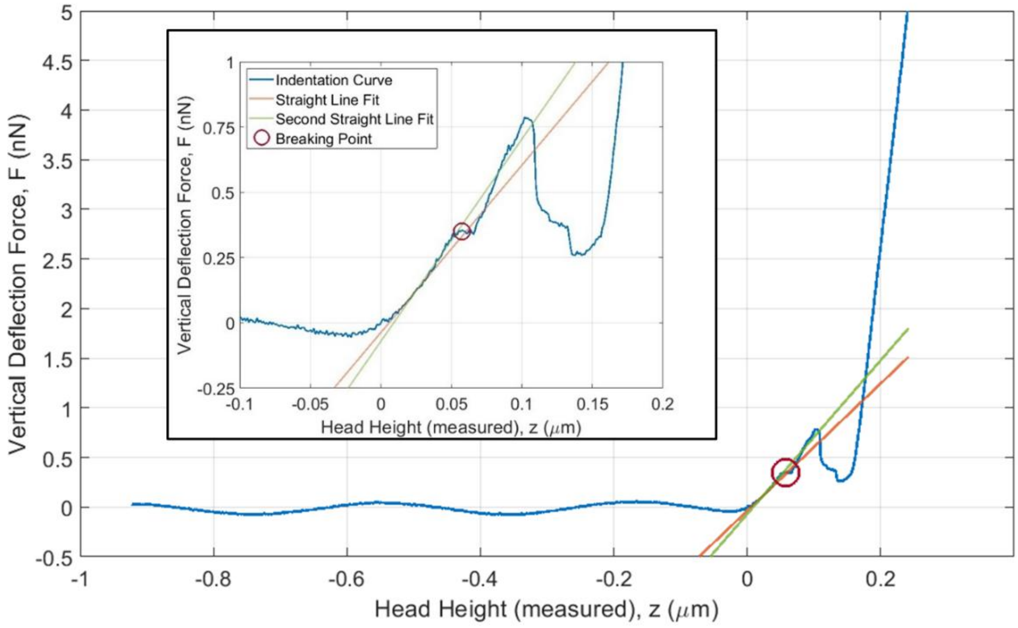


Figure S2. The vertical deflection force and AFM head height for a specific nanoindentation experiment indicating the first slope fit, breaking point and the second slope fit. Insert is the close-up view of the region of interest.

Table S1. Seed distance, number of elements, and predicted force value for the mesh convergence study.

| Seed distance | No. of Elements | Max Force Value |
|---|---|---|
| 0,8 | 278976 | 0,199849 |
| 1 | 235776 | 0,199644 |
| 1,2 | 217344 | 0,19283 |
| 1,4 | 200448 | 0,198344 |
| 1,6 | 186624 | 0,191301 |
| 1,8 | 174144 | 0,191318 |
| 2 | 165312 | 0,197224 |
| 2,3 | 153600 | 0,185954 |
| 2,5 | 151488 | 0,184576 |
| 2,5 | 96768 | 0,180902 |
| 2,6 | 94066 | 0,181783 |
| 2,7 | 73956 | 0,169117 |
| 2,9 | 63504 | 0,155757 |
| 3 | 57600 | 0,158873 |
| 3,25 | 41790 | 0,161612 |
| 3,5 | 37430 | 0,164510 |
| 3,7 | 33310 | 0,164829 |
| 4 | 21600 | 0,075344 |
| 4,25 | 20632 | 0,076697 |
| 4,5 | 17912 | 0,0807073 |
| 5,5 | 8712 | 0,0949015 |
| 8,5 | 2236 | 0,129554 |
| 12 | 1200 | 0,156753 |
| 18 | 696 | 0,174782 |
| 25 | 256 | 0,207845 |
| 50 | 80 | 0,219744 |
